# Mechanical Stimuli such as Shear Stress and Piezo1 Stimulation Generate Red Blood Cell Extracellular Vesicles

**DOI:** 10.1101/2022.10.04.510887

**Authors:** Gurneet S. Sangha, Callie M. Weber, Ryan M. Sapp, Saini Setua, Kiruphagaran Thangaraju, Morgan Pettebone, Allan Doctor, Paul W. Buehler, Alisa M. Clyne

## Abstract

Circulating red blood cell extracellular vesicles (RBC-EVs) are a promising biomarker for vascular health. However, generating, isolating, and characterizing physiologically relevant RBC-EVs with sufficient yield and purity for biological studies is non-trivial. Here, we present and rigorously characterize an *in vitro* model to mimic RBC-EV production during shear stress via mechanosensitive piezo1 ion channel stimulation. We optimize our RBC-EV isolation protocol to minimize hemolysis, maximize RBC-EV yield and purity, and improve the ease of EV characterization. RBC-EV purity was measured by quantifying protein (e.g., particles/*μ*g), large particle (e.g., protein aggregates), and platelet EV contamination. This study compared RBC-EV isolation performance using membrane-based affinity (e.g., exoEasy), ultrafiltration (e.g., Amicon Ultra-15), and ultracentrifugation, with and without size exclusion chromatography purification. We found that treating 6% hematocrit with 10 *μ*M piezo1-agonist yoda1 for 30 minutes and isolating RBC-EVs using ultracentrifugation minimized RBC hemolysis and maximized RBC-EV yield (~10^12^ particles/mL) and purity, provided the most consistent RBC-EV preparations, and improved ease of RBC-EV characterization. Our pressure myography experiments suggest that co-isolated protein contaminants, but not piezo1 RBC-EVs, induce rapid mouse carotid artery vasodilation. These results underscore the importance of characterizing EV purity for biological experiments. The standardized methods outlined here enable mechanistic studies of how RBC-EVs generated in physiological flow affect vascular response.

## Introduction

Red blood cells (RBCs) are exposed to both chemical and physical stresses as they circulate throughout the body. Perhaps in response to these stresses, RBCs lose 20% of their volume during their 120-day lifecycle.^1,2^ Some RBC volume loss comes in the form of extracellular vesicles (RBC-EVs), defined as lipid-bound particles 40-1000 nm in diameter that transport signaling molecules (e.g., proteins, RNA) to other cells.^3,4^ RBC-EVs generated in stress associated with pathological conditions, such as obstructive sleep apnea,^5^ sickle cell disease,^6–8^ or blood storage^9,10^ exacerbate endothelial dysfunction, the initial hallmark of vascular disease. These atherogenic RBC-EV effects are partly attributed to hemoglobin, which can decrease endothelial nitric oxide synthase (eNOS) phosphorylation, scavenge nitric oxide (NO), and increase oxidative stress.^11,12^ However, RBC-EVs are likely also generated during physiological mechanical stress, including shear stress, as RBCs (8-10 *μ*m diameter) squeeze through capillaries (2-4 *μ*m diameter). It remains unclear how circulating RBC-EVs generated under physiological shear stress affect endothelial function and subsequent vascular health.

Shear stress may trigger RBC-EV production by stimulating piezo1, a mechanosensitive Ca^2+^ ion channel.^13,14,15^ A seminal study by Larsen *et al.* showed that shear stress enhances RBC Ca^2+^ permeability.^16^ Danielczok *et al.* later confirmed that RBCs squeezing through capillaries transiently increase intracellular Ca^2+^ through piezo1 activation.^17^ RBC Ca^2+^ uptake activates scramblase, a transmembrane protein that translocates RBC phosphatidylserine from the inner to outer plasma membrane leaflet to trigger RBC-EV release.^18–20^ Therefore, piezo1 activation may also generate RBC-EVs through similar Ca^2+^-dependent mechanisms. Taken together, we hypothesize that shear stress and mechanosensitive piezo1 stimulation increases RBC Ca^2+^ influx to trigger RBC-EV production.

Investigating RBC-EVs and their impact on vascular function is challenging due to the variety of EV production and isolation methods. RBC-EVs can be isolated from human blood; however, these samples contain EVs from multiple cell types, as well as non-EV contaminants such as large protein aggregates and lipoproteins.^21,22^ Cleaner RBC-EVs can be produced *in vitro* by treating RBCs with *tert*-butyl hydroperoxide to induce oxidative stress,^23^ oxygen cycling to mimic RBC hypoxia,^5^ Ca^2+^ ionophore A23187 to model eryptosis,^24^ needle shearing or high-pressure extrusion to simulate shear stress,^6,25^ or long term storage to produce RBC-EVs relevant in blood transfusions.^9,10^ RBC-EVs can then be isolated and concentrated using membrane-based affinity, ultrafiltration, or ultracentrifugation. Each of these techniques for producing and isolating EVs can impact RBC-EV concentration, contents, and purity, and therefore the biological function of these RBC-EVs.

EVs have emerged as a potential biomarker for vascular disease, as they may serve as conduits for intercellular signaling throughout the body. RBC-EVs generated during shear stress are of particular interest because they may significantly contribute to the circulating EV pool under non-pathological conditions; however, the lack of standards for mechanically-stimulated RBC-EVs hinders biological studies. In this paper, we develop methods to produce and isolate RBC-EVs via mechanical stimulation with sufficient concentration and purity for mechanistic studies. We show that both shear stress and piezo1 stimulation trigger RBC-EV release. RBC piezo1 stimulation and RBC-EV isolation protocols are fine-tuned to maximize RBC-EV production and minimize RBC hemolysis and contaminants. Finally, we use pressure myography to highlight how RBC-EV purity can affect biological studies. The methods outlined are designed to be implemented by researchers interested in studying shear stress-induced piezo1 RBC-EVs.

## Results

### RBC-EVs circulate in healthy humans and are released by RBCs under shear stress

We performed flow cytometry to identify EVs isolated from healthy human plasma based on CD63 and CD81 expression (**Fig. 1a**). The primary cellular sources for EVs in healthy human plasma were identified as platelets (CD41+), RBCs (CD235a+), and endothelial cells (CD31+; **Fig. 1b**). Next, we studied whether RBCs exposed to shear stress produce RBC-EVs. We needle-sheared RBCs and quantified particle concentration and size in the RBC supernatant using TRPS. Needle-sheared RBCs generated 3.6×10^9^ ± 2.7×10^9^ particles/mL, with particle sizes ranging from 55.3 - 128.7 nm (**Fig. 1c**). The particle rate from needle-sheared RBCs was 29-fold higher compared to RBCs in static conditions (**Fig. 1d**; *p*<0.05). To demonstrate that these particles are EVs, we lysed the particles with Triton X-100 and counted the remaining particles. Indeed, particle rate decreased 10-fold when needle sheared samples were treated with 1% Triton X-100 (**Fig. 1d**). Western blot confirmed that the shear stress-induced RBC particles contained cytoplasmic EV markers ALIX and TSG101 and surface EV marker CD63 (**Fig. 1e**). RBC particles generated by shear stress also contained more RBC marker stomatin than particles isolated from RBCs that did not undergo shear stress (**Fig. 1e**). Finally, TEM demonstrated that the shear stress-generated RBC particles resemble EVs that were ~100 nm in diameter (**Fig. 1f**).

**Fig. 1.**
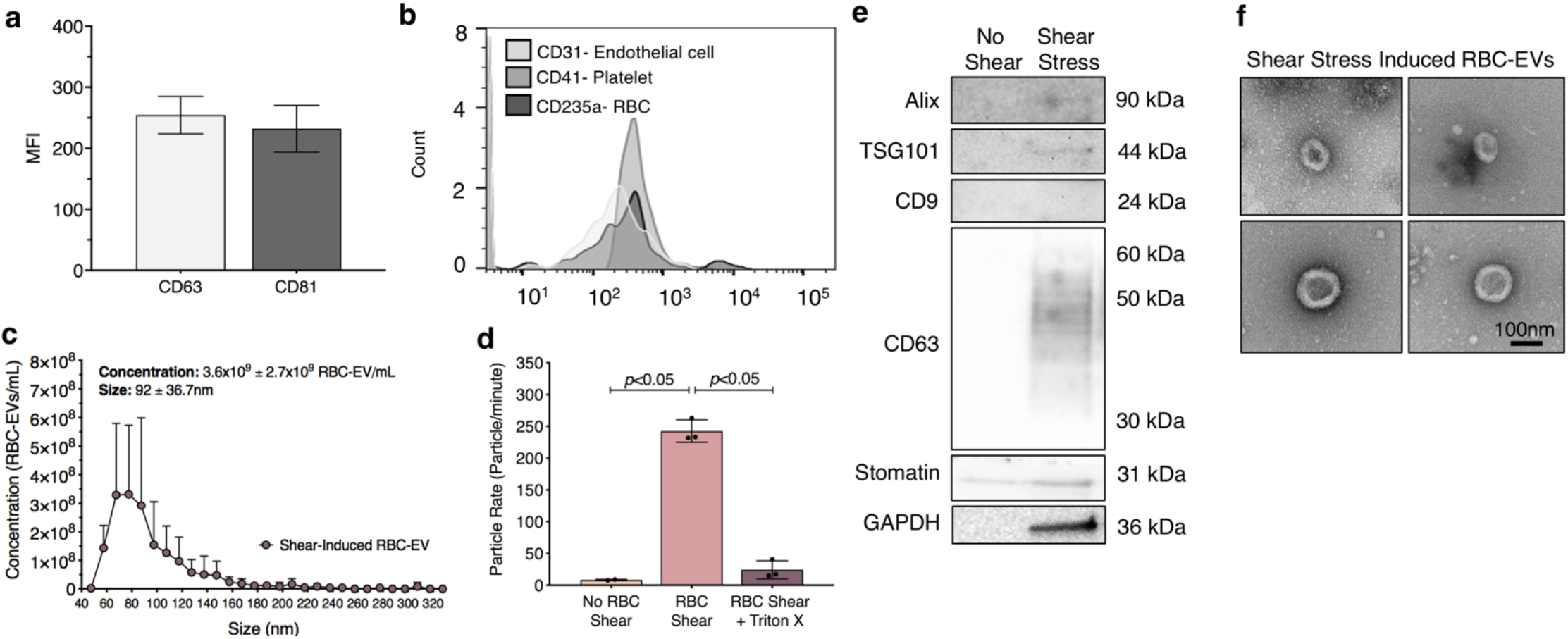
Circulating human RBC-EV detection and shear-induced RBC-EV characterization. **a.** Quantified flow cytometry mean fluorescence intensity (MFI) of plasma EV samples (n=6) for EV markers CD63 and CD81. **b.** Representative flow cytometry histogram of plasma EVs for RBC (CD235a), platelet (CD41), and endothelial cells (CD31) markers. **c.** Size histogram of particles isolated from needle sheared RBCs using Amicon Ultra-15 centrifuge units. **d.** Particle rate of samples isolated from RBCs in static conditions, RBC exposed to shear stress, and RBC exposed to shear stress with isolated particles treated with 1% Triton X-100. Particle rate was quantified by measuring the number of particles that pass through the TRPS nanopore over one minute. **e.** Western blot of needle-sheared RBC-EVs for cytoplasmic EV marker ALIX and TSG101, surface EV markers CD9 and CD63, RBC-specific marker stomatin, and housekeeper GAPDH. Each Western blot shows one representative comparison from the full blot. Western blots were performed using n=1 biological replicate and repeated three times using RBCs from three different blood donors **f.** TEM images of RBC-EVs generated through needle shearing.

### Piezo1 stimulation maintains RBC viability and decreases RBC hemolysis compared to Ca^2+^ ionophore treated RBCs

Shearing RBCs through a needle is labor intensive and time-consuming, while other methods to shear RBCs require specialized equipment that are not accessible to all investigators. Therefore, we investigated how chemical stimulation of the mechanosensitive ion channel Piezo1 via the agonist yoda1 affects RBCs. We compared yoda1-treated RBCs to Ca^2+^ ionophore-treated RBCs, as Ca^2+^ ionophore is a standard method for RBC-EV generation. Untreated RBCs remained mostly viable after 24 hours incubation at 37°C, as determined by calcein labeling (**Fig. 2a**). Approximately 80% of RBCs were viable after 24 hours of treatment with 0.28% DMSO (vehicle control) or 100 *μ*m yoda1. RBC viability significantly decreased to 64±17% with 100 *μ*m Ca^2+^ ionophore treatment (*p*<0.05 compared to freshly isolated RBCs; **Fig. 2b**). Ca^2+^ ionophore treated RBCs also experienced greater hemolysis (**Fig. 2c**) and produced more cell debris (**Supplementary Fig. 1**) than yoda1 treated RBCs.

**Fig. 2.**
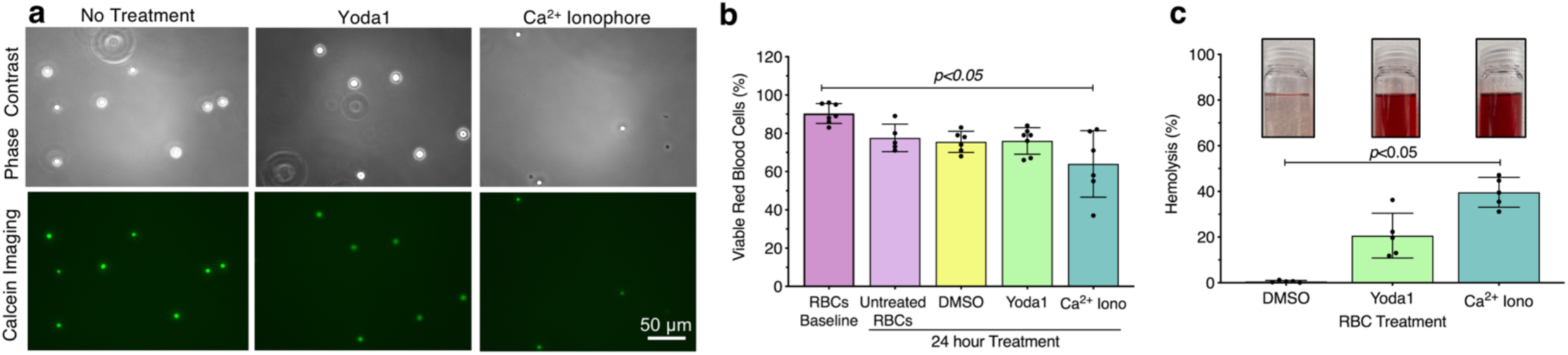
RBC viability, hemolysis, and eNOS phosphorylation measurements after piezo1 stimulation and Ca^2+^ ionophore treatment. **a.** Representative calcein-labeled RBC fluorescence microscopy images, with **b.** quantification to assess RBC viability **c.** RBC hemolysis due to 0.28% DMSO, 100 *μ*M piezo1-agonist yoda1, and 100 *μ*M Ca^2+^ ionophore treatment for 24 hours, as measured by hemoglobin absorbance at 541nm in the RBC supernatant. Insets show representative images of RBC supernatant after treatment.

### Piezo1 stimulation and Ca^2+^ ionophore treatment yield similar RBC-EV quantity and purity

We then compared the size and concentration of RBC-EV produced via yoda1 and Ca^2+^ ionophore RBC treatment. Yoda1 treatment produced 2.7×10^12^ ± 2.8×10^12^ RBC-EVs/mL with a diameter of 125 ± 45 nm (**Fig. 3a,b**). Ca^2+^ ionophore treatment produced 3.6×10^12^± 6.2×10^12^ RBC-EVs/mL with a similar size (122 ± 41 nm diameter; **Fig. 3a,b**). DMSO vehicle control produced 1000X fewer EVs (6.3×10^9^± 3.7×10^9^ particle/mL) with a smaller size (87 ± 28 nm diameter; **Fig. 3a,b**). Piezo1 and Ca^2+^ ionophore RBC-EVs had similar purity, as determined by the particle/total protein ratio (**Fig. 3c**). Both piezo1 and Ca^2+^ ionophore RBC-EVs contained cytoplasmic EV markers ALIX and TSG101 and surface EV markers CD9 and CD63, along with RBC specific marker stomatin (**Fig. 3d**). ALIX quantity was consistently greater in Ca^2+^ ionophore as compared to piezo1 RBC-EVs (*p*>0.05; **Supplementary Fig. 2**). TEM confirmed that piezo1 and Ca^2+^ ionophore RBC-EVs were similar in size and morphology (**Fig. 3e**).

**Fig. 3.**
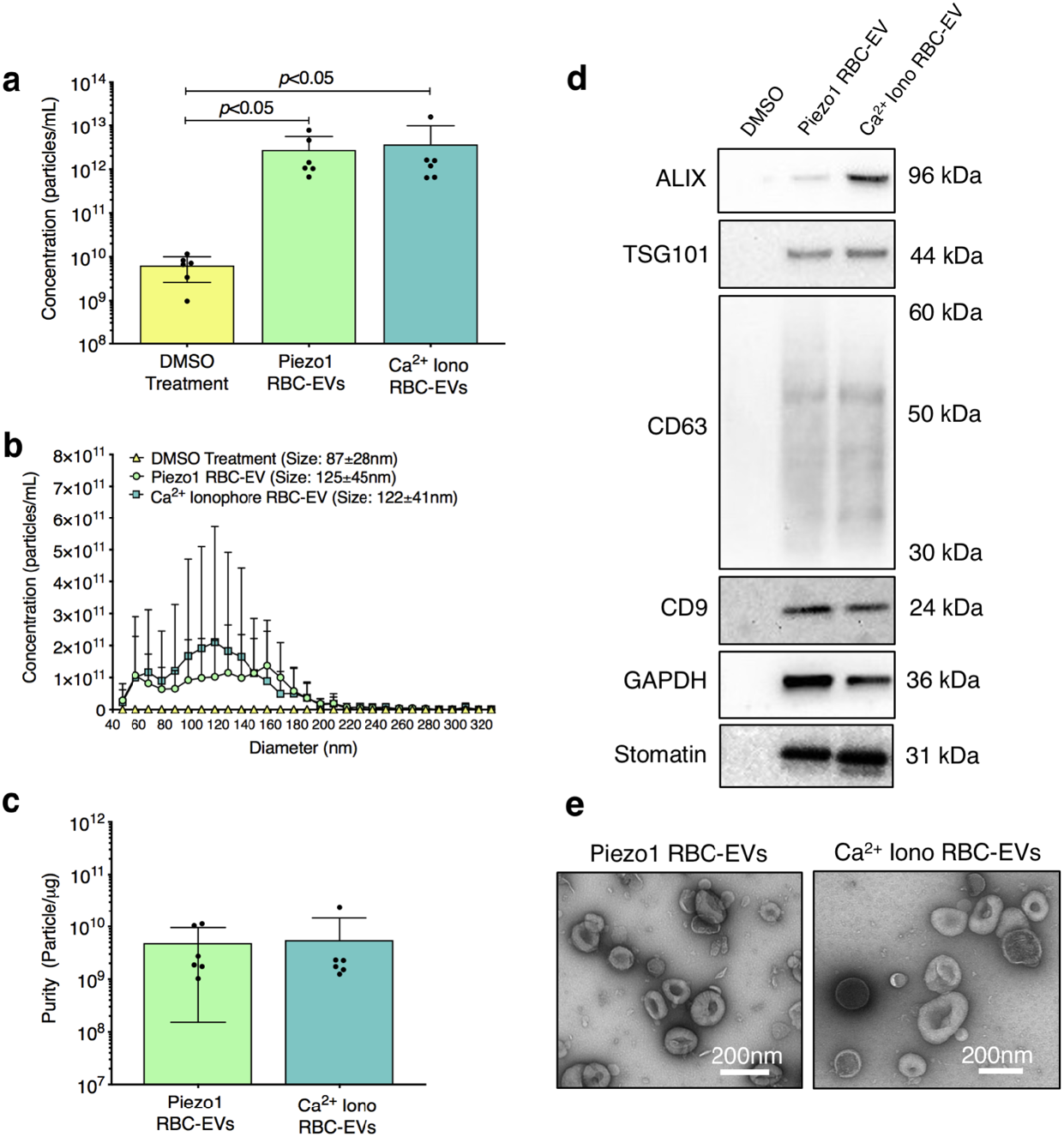
Piezo1 and Ca^2+^ ionophore RBC-EV concentration, size, EV marker, and morphology characterization. **a.** Particle concentration in RBC supernatant after 0.28% DMSO, 100 *μ*M yoda1, and 100 *μ*M Ca^2+^ ionophore treatment for 24 hours, as measured by TRPS **b**. Size histogram of particles isolated from RBC supernatant after DMSO, yoda1, and Ca^2+^ ionophore treatment, as measured by TRPS **c.** Piezo1 and Ca^2+^ ionophore RBC-EV purity, as measured by calculating particles/*μ*g total protein. **d.** Representative Western blots of cytoplasmic EV markers ALIX and TSG101, surface EV markers CD9 and CD63, RBC-specific marker stomatin, and housekeeper GAPDH. Western blots were performed using n=3 biological replicates and repeated two times. **e**. Representative TEM images of piezo1 and Ca^2+^ ionophore RBC-EVs.

### 6% hematocrit treated with 10 *μ*M yoda1 for 30 minutes minimized hemolysis and maximized RBC-EV concentration and purity

We next determined the effect of hematocrit, yoda1 incubation time, and yoda1 concentration on piezo1 RBC-EV concentration, size, and purity. 1% hematocrit treated with 100 *μ*M yoda1 for 2 hours generated 2.1×10^11^±7.7×10^10^ RBC-EV/mL. The RBC-EV concentration increased with hematocrit, with 3% hematocrit producing 6.7×10^11^±3.1×10^11^ RBC-EV/mL, 6% hematocrit producing 4.7×10^12^±1.9×10^12^ RBC-EV/mL, and 18% hematocrit producing 1×10^13^± 5.7×10^12^ RBC-EV/mL (**Fig. 4a,b**). While RBC-EV size was consistent across all hematocrits, RBC-EVs produced using 6% and 18% hematocrit had greater purity than RBC-EVs produced using 1% (*p*>0.05) and 3% hematocrit (*p*<0.05; **Fig. 4c**). We were also able to robustly detect cytoplasmic EV markers ALIX and TSG101, surface EV markers CD9 and CD63, and RBC-EV marker stomatin by Western blot when using 6% and 18% hematocrit (**Fig. 4d**). We did not detect apoptotic body marker Calnexin in RBC-EVs generating using up to 18% hematocrit. We therefore used 6% hematocrit for the yoda1 incubation time experiments. RBC-EVs ranging from 85 to 153 nm diameter were generated within 30 minutes (**Fig. 4e,f**). There was no difference in RBC-EV concentration, average size, or purity after 0.5, 1, 2, or 24 hours of yoda1 treatment (**Fig. 4e,f**). However, 24 hours of yoda1 treatment generated three distinct RBC-EV size peaks at approximately 70, 110, and 150 nm diameter, while RBCs treated with yoda1 for 2 hours or less generated one RBC-EV peak at 110 nm diameter (**Fig. 4f**). We therefore used 6% hematocrit and 30 minutes incubation time for the yoda1 concentration experiments. RBCs treated with 10 *μ*M yoda1 produced significantly more RBC-EVs than RBCs treated with 1 *μ*M yoda1 (**Fig 4h,i**). There was no difference in RBC-EV purity with 1, 10, 100 *μ*M yoda1 or 10 *μ*M calcium ionophore treatment (**Fig. 4j**). Western blots of RBC-EV generated at different hematocrit, yoda1 incubation time, and yoda1 concentrations are in **Supplementary Fig. 3.** The improved RBC-EV production method (6% hematocrit exposed to 30 minutes of 10 *μ*M yoda1) induced significantly less hemolysis than the original RBC-EV production method (6% hematocrit exposed to 24 hours of 100 *μ*M yoda1; **Fig. 4k**).

**Fig. 4.**
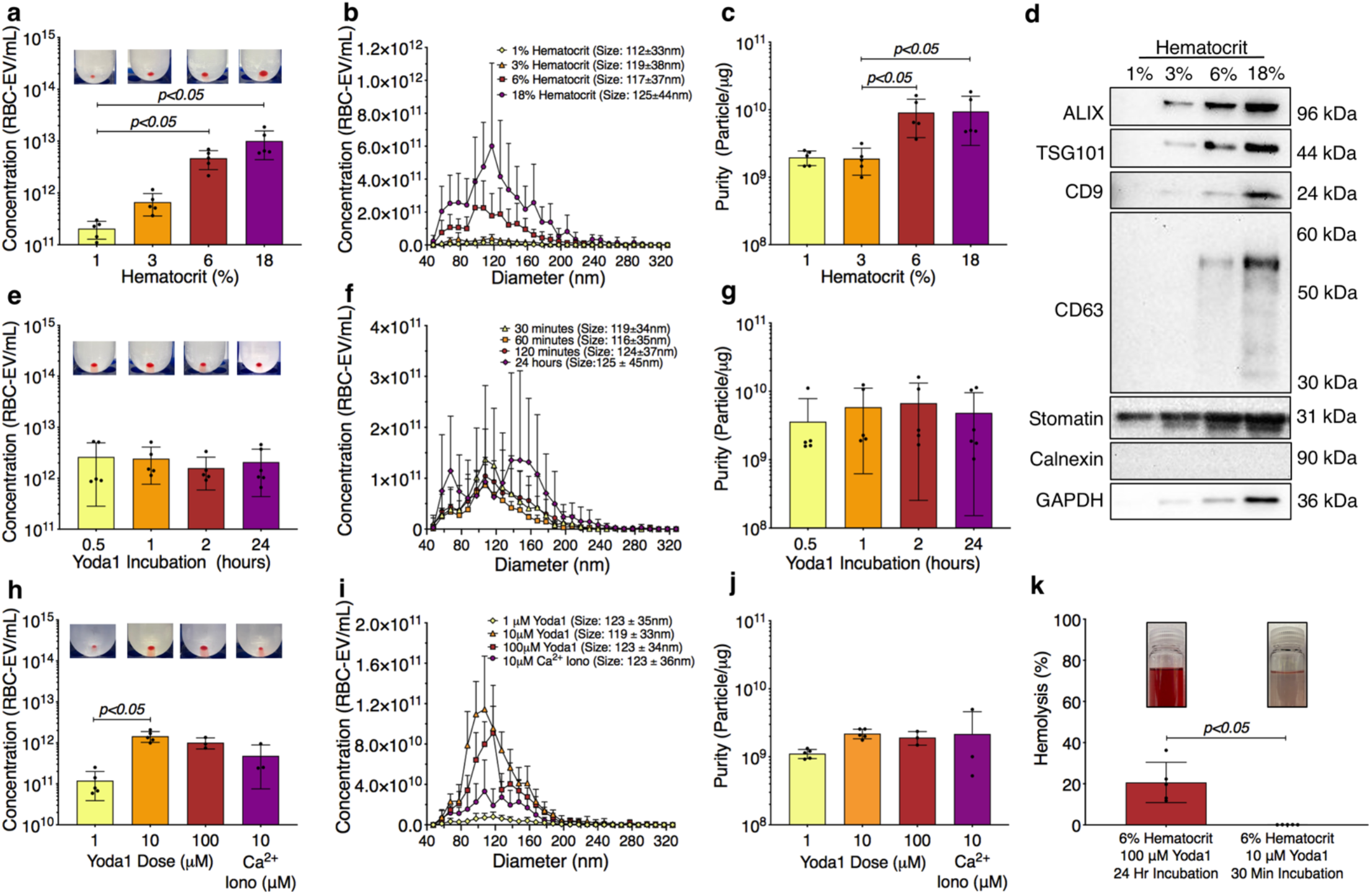
Optimization of hematocrit, yoda1 incubation time, and yoda1 dose to minimize hemolysis and maximize RBC-EV yield and purity. **a.** RBC-EV concentration (TRPS), **b.** size histogram (TRPS), and **c.** protein contamination (particle/*μ*g protein) for 1, 3, 6, and 18% hematocrit treated with 100 *μ*M yoda1 for 2 hours. **d.** Representative western blot of cytoplasmic EV markers ALIX and TSG101, surface EV markers CD9 and CD63, RBC-specific marker stomatin, apoptotic body marker Calnexin, and housekeeper GAPDH. Western blots were performed using n=3 biological replicates and repeated two times. **e.** RBC-EV concentration (TRPS), **f.** size histogram (TRPS), and **g.** purity (particle/*μ*g protein) for 6% hematocrit treated with 100 *μ*M yoda1 for 0.5, 1, 2, and 24 hours. **h.** RBC-EV concentration (TRPS) **i.** size histogram (TRPS), and **j.** purity (particle/*μ*g protein) for 6% hematocrit treated with 1, 10, 100 *μ*M yoda1 or 10 *μ*M Ca^2+^ ionophore for 30 minutes. Inset in panels a, e, and h shows RBC-EV pellet after second ultracentrifugation at 100,000xg. **k.** Hemolysis using original treatment (6% hematocrit, 100 *μ*M yoda1, 24 hour treatment) and optimized treatment (6% hematocrit, 10 *μ*M yoda1, 30 minute treatment) for RBC-EV production. Inset in panel k shows representative RBC supernatant to show hemolysis.

### Ultracentrifugation and Amicon Ultra-15 centrifuge units enabled reliable RBC-EV isolation and characterization

We next compared the yield and purity of RBC-EVs isolated using membrane-based affinity binding (exoEasy), ultrafiltration (Amicon Ultra-15), and ultracentrifugation. When RBC-EVs from each isolation method were purified using size exclusion chromatography (SEC), which eluted the RBC-EVs into six, 500 *μ*L fraction. Protein concentration measurements showed that RBC-EVs were in the first three SEC fractions from ultrafiltration and ultracentrifugation samples (**Fig. 5a**). We were unable to identify RBC-EVs in any SEC fractions for samples isolated using exoEasy. Ultracentrifugation without SEC generated the most concentrated RBC-EV (8.2×10^10^±2.5×10^10^ particle/mL/mL) compared to exoEasy without SEC (*p*<0.05; 9.4×10^9^±3.3×10^9^ particle/mL/mL) and Amicon without SEC (4.1×10^11^±3.5×10^10^ particle/mL/mL). SEC consistently decreased RBC-EV concentration (3.7×10^11^ ± 6.9×10^10^ particle/mL for ultracentrifugation with SEC, 2.3×10^11^±1.2×10^10^ particle/mL for Amicon Ultra-15 with SEC; **Fig. 5b**). RBC-EV isolation without SEC yielded particles 108.2 ± 45.9 nm in diameter using exoEasy, 115.2 ± 44.9 nm in diameter using Amicon Ultra-15, and 103.6 ± 36.1nm in diameter using ultracentrifugation (**Fig. 5c**). RBC-EV isolation with SEC did not significantly alter RBC-EV size (**Fig. 5d**). All cytoplasmic and surface EV markers were detected by Western blot in RBC-EVs isolated using Amicon Ultra-15 and ultracentrifugation; however, EV markers were not detected from particles isolated using exoEasy (**Fig. 5e**). TEM verified our TRPS and Western blot results, revealing sparse EVs in samples isolated using exoEasy, more concentrated EVs in samples isolated using Amicon Ultra-15, and most concentrated EVs in samples isolated using ultracentrifugation (**Fig. 5f**).

**Fig. 5.**
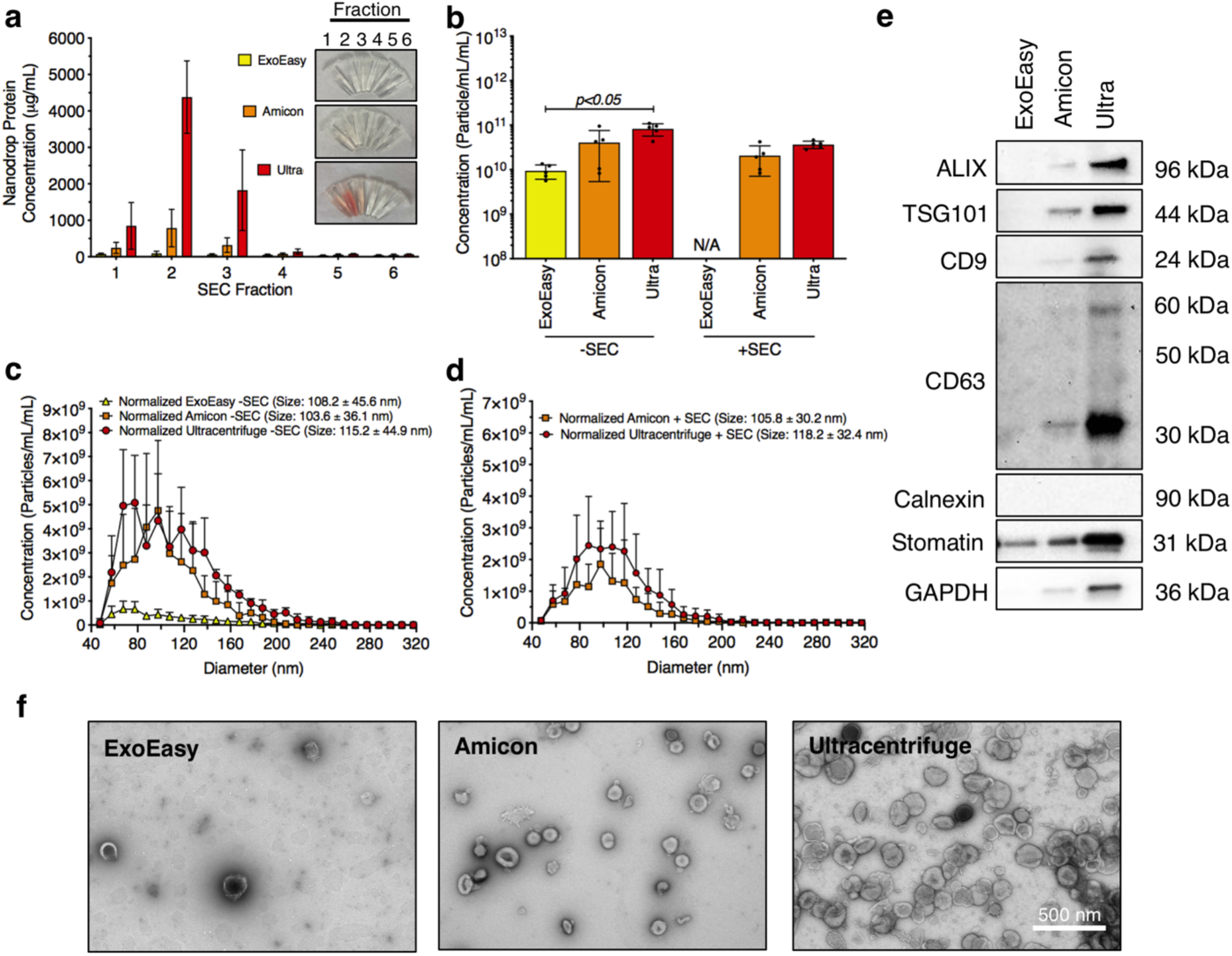
Piezo1 RBC-EV quantification and characterization using exoEasy, Amicon Ultra-15, and ultracentrifugation isolation methods, with and without SEC purification. **a.** Nanodrop protein measurements of SEC fractions from RBC-EVs isolated using exoEasy, Amicon Ultra-15, and ultracentrifugation. **b.** Concentration of RBC-EVs isolated using exoEasy, Amicon Ultra-15, and ultracentrifugation with and without SEC purification. Concentration was normalized to the starting RBC supernatant volume used for the RBC-EV isolation. **c.** Size histogram of RBC-EVs isolated using exoEasy, Amicon Ultra-15, and ultracentrifugation *without* SEC purification. **d.** Size histogram of RBC-EVs isolated using exoEasy, Amicon Ultra-15, and ultracentrifugation *with* SEC purification. **e.** Western blots of cytoplasmic EV markers ALIX and TSG101, surface EV markers CD9 and CD63, RBC-specific marker stomatin, apoptotic body marker calnexin, and housekeeper GAPDH. Each Western blot shows one representative comparison. Western blots were performed using n=3 biological replicates and repeated twice. **f.** TEM of RBC-EVs isolated using exoEasy, Amicon Ultra-15, and ultracentrifugation. RBCs (6% hematocrit) were treated with 10 *μ*M yoda1 for 30 minutes.

### RBC-EV isolation by ultracentrifugation enabled the highest yield and the least protein contamination

We then compared RBC-EV isolation methods for yield and purity by measuring RBC-EV protein contamination. Particles isolated using exoEasy were less pure (8.2×10^8^±3.8×10^8^ particles/*μ*g protein) than RBC-EVs isolated by ultracentrifugation 2.8×10^9^±7.3×10^8^ particles/*μ*g; *p*<0.05; **Fig. 6a**). SEC did not improve purity in RBC-EVs isolated using Amicon Ultra-15 or ultracentrifugation (**Fig. 6a**). Protein aggregates were measured using EV/non-EV particle rate ratio, calculated by quantifying sample particle rate before and after lysing RBC-EVs using Triton-X 100 surfactant. EV/non-EV particle rate ratio suggests that there is no statistically significant difference in protein aggregate contamination amongst exoEasy, Amicon Ultra-15, and ultracentrifugation RBC-EV isolation methods (**Fig. 6b**). SEC decreased RBC-EV protein aggregates 2.3-fold after ultracentrifugation and 3.8-fold after Amicon Ultra-15 isolation, but the results were variable amongst the biological replicates (**Fig. 6b**). Finally, RBC-EVs isolated by ultracentrifugation contained significantly less platelet EV contamination (CD41+) compared to particles isolated using exoEasy, as measured by flow cytometry (**Fig. 6c,d**). SEC did not affect platelet contamination in EV samples isolated using Amicon Ultra-15 or ultracentrifugation (**Fig 6c,d**).

**Fig. 6.**
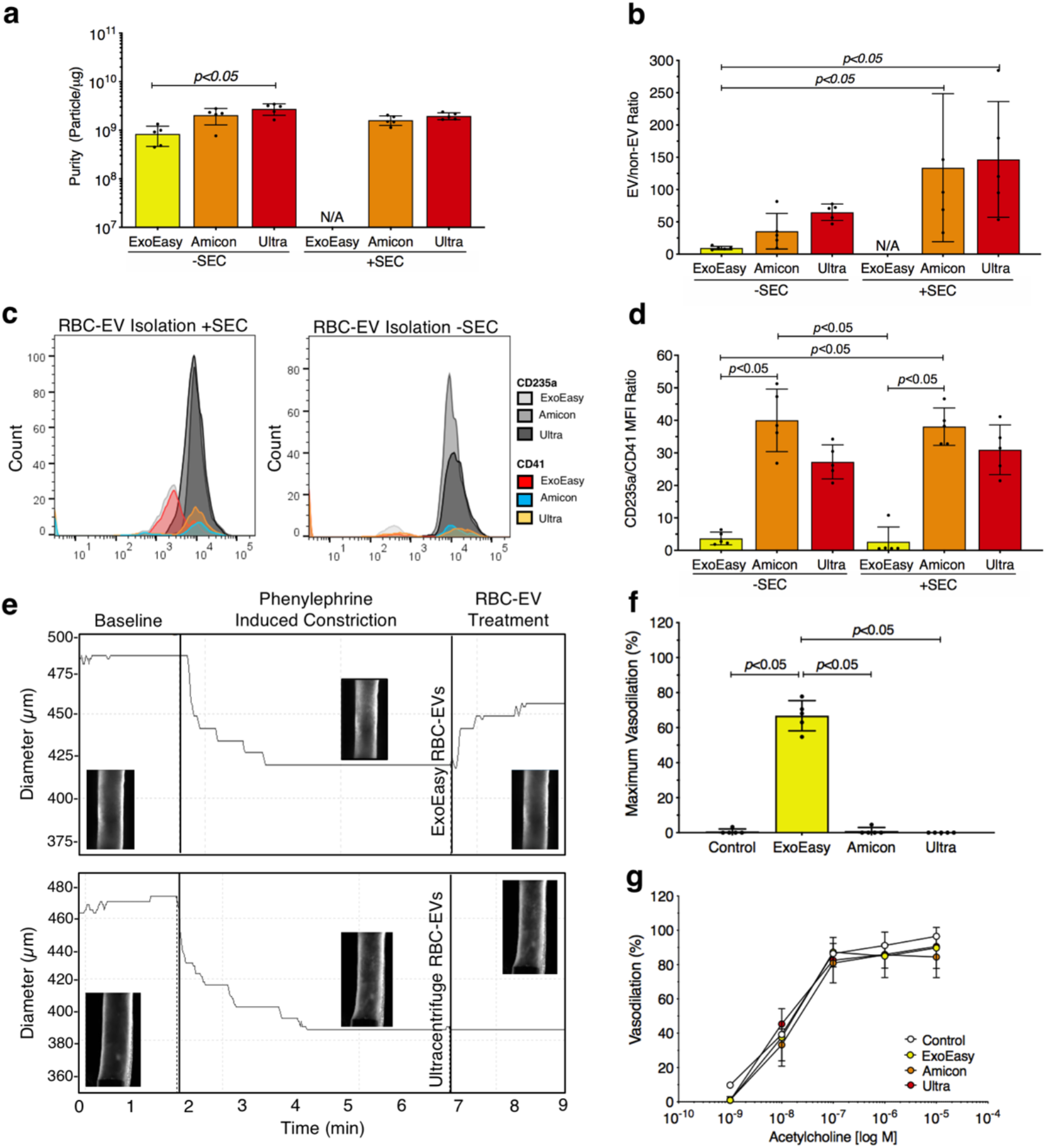
Characterization of piezo1 RBC-EV protein, protein aggregate, and platelet EV contamination and their effects on mouse carotid artery function. **a.** Purity of RBC-EVs isolated using exoEasy, Amicon Ultra-15, and ultracentrifugation, measured by calculating particle/*μ*g protein. **b.** Protein aggregates in RBC-EVs isolated using exoEasy, Amicon Ultra-15, and ultracentrifugation. EV/non-EV ratio was calculated by quantifying sample qNano particle rate before and after lysing RBC-EVs using Triton-X 100 surfactant. **c.** Representative flow cytometry histogram for RBC (CD235a+) and platelet (CD41+ markers). **d.** Platelet RBC-EV contamination for RBC-EVs isolated using exoEasy, Amicon Ultra-15, and ultracentrifugation with and without SEC, quantified by the RBC(CD235a+):Platelet(CD41+) ratio from flow cytometry. **e.** Representative pressure myography data of phenylephrine constricted mouse carotid arteries treated with RBC-EVs. **f.** Maximum mouse carotid artery vasodilation induced by RBC-EV treatment. **g.** Endothelial-dependent mouse carotid artery vasodilation after 5 minutes RBC-EV treatment, measured using acetylcholine dose response. Controls include buffer XE used to resuspend exoEasy isolated RBC-EVs and 2xPBS used to resuspend Amicon Ultra-15 and ultracentrifuge isolated RBC-EVs.

### Protein contaminants in isolated RBC-EV triggered carotid artery vasodilation, while piezo1 RBC-EVs did not inhibit endothelial dependent vasodilation

**T**o determine if RBC-EV purity impacted functional assays, we studied how RBC-EVs isolated using exoEasy, Amicon Ultra-15, and ultracentrifugation affected mouse carotid artery vasodilation. Pre-constricted mouse carotid arteries treated with exoEasy isolated RBC-EV rapidly vasodilated 66 ± 8.6%, while Amicon Ultra-15 and ultracentrifuge isolated RBC-EVs generally did not cause vasodilation (**Fig. 6e,f**). Carotid arteries treated with RBC-EV solvents 2xPBS and buffer XE did not cause vasodilation. Endothelial dependent vasodilation, as measured by acetylcholine dose response in mouse carotid arteries, did not significantly change with RBC-EV or vehicle control treatment (**Fig. 6g**). Untreated carotid arteries maximally vasodilated 96.5 ± 3.2%, while exoEasy, Amicon Ultra-15, and ultracentrifugation piezo1 RBC-EV treated carotid arteries maximally vasodilated 89.7 ± 12.0%, 84.4 ± 12.0%, and 90.6 ± 11.7% respectively (**Fig. 6g**).

### RBC-EVs isolated from fresh and frozen RBC supernatant had similar yield, size, and purity

Finally, we investigated how storage affects RBC-EV concentration, size, and purity. First, we isolated RBC-EVs from RBC supernatant that was either fresh or had undergone one freeze-thaw cycle (-80°C to 25°C). We found no significant differences in EV concentration (**Supplementary Fig. 4a**), purity (**Supplementary Fig. 4b**), and size (**Supplementary Fig. 4c**) between RBC-EVs isolated using fresh or frozen RBC supernatant. We then subjected RBC-EVs isolated from fresh RBC supernatant to one, two, or three freeze-thaw cycles. RBC-EVs that underwent 1 freeze-thaw cycle contained 2.4×10^12^ ± 2.3×10^11^ RBC-EV/mL at size 109 ± 31 nm, which was similar to the concentration prior to the freeze-thaw cycle. However, RBC-EV concentration decreased to 5.0×10^11^ ± 3.1×10^10^ RBC-EV/mL and 5.8×10^11^ ± 8.3×10^10^ RBC-EV/mL after the second and third freeze-thaw cycles (**Supplementary Fig. 4d,f**). RBC-EV size also significantly decreased to 73.8 ± 16.5 nm and 83 ± 14.8 nm after the second and third freeze-thaw cycles (*p*<0.05; **Supplementary Fig. 4e,f**). Moreover, RBC-EVs that underwent one freeze thaw cycle and then were stored at 4°C gradually decreased in concentration (2.4×10^12^ ± 2.3×10^11^ at baseline to 1.0×10^12^ ± 1.5×10^11^ at day 7; **Supplementary Fig. 4g,i**) and rapidly decreased in size (112.4 ± 31.8 nm diameter to 77.8 ± 13.9 nm over 7 days; **Supplementary Fig. 4h,i**). RBC-EV size versus concentration histogram suggests that smaller RBC-EVs remain more stable during multiple freeze thaw cycles and 4°C storage (**Supplementary Fig. 4f,i**).

## Discussion

RBC-EVs generated during shear stress are of particular interest because they may significantly contribute to the circulating EV pool under non-pathological conditions; however, the lack of standards for mechanically-stimulated RBC-EVs hinders mechanistic studies. In this paper, we develop methods to produce and isolate RBC-EVs via mechanical stimulation with sufficient concentration and purity for mechanistic studies. We show that both shear stress and piezo1 stimulation trigger RBC-EV release. RBC piezo1 stimulation and RBC-EV isolation protocols are fine-tuned to maximize RBC-EV production and minimize RBC hemolysis and contaminants. Finally, we use pressure myography to highlight how RBC-EV purity can affect biological studies. The methods outlined in the manuscript are designed to be implemented by researchers interested in RBC-EVs and further our understanding of shear stress-induced piezo1 RBC-EVs.

RBC-EVs circulate in healthy humans, suggesting a non-pathological function for RBC-EVs that may regulate vascular health. Indeed, our flow cytometry experiments show that plasma contains a subfraction of EVs expressing the RBC marker CD235a. Nemkov *et al*. also showed that healthy individuals have circulating RBC-EVs, detecting <1 *μ*m diameter particles with both RBC marker CD235a and EV marker phosphatidylserine by flow cytometry.^26^ Moreover, they found that short-term exercise cycling may increase RBC-EV shedding,^26^ potentially due to augmented capillary perfusion that increases RBC shear stress exposure. We and others similarly found that needle shearing RBCs (shear stress ~120 dynes/cm^2^) release EVs.^6^ RBCs begin to deform around 5-20 dynes/cm^2^ and rupture around 1500 dynes/cm^2^.^27,28^ Therefore, needle shearing RBCs should induce EV production via RBC deformation and not hemolysis. We also observed <1% hemolysis with RBC needle shearing, suggesting that RBC-EVs were not formed by rupture. It is unclear, however, how shear stress triggers RBCs to release RBC-EVs.

Our data show that stimulation of mechanosensitive piezo1 causes RBCs to release RBC-EVs. It is well established that piezo1 opens and increases RBC Ca^2+^ uptake in response to mechanosensitive stimuli, such as shear stress. Therefore, it is likely that hemodynamic stimuli during physiological blood flows triggers piezo1 stimulation and subsequent RBC-EV release. However, increasing intracellular RBC Ca^2+^ concentration using Ca^2+^ ionophore is typically used to model red blood cell death or eryptosis. Indeed, we found that Ca^2+^ ionophore treatment significantly increased RBC death quantified using calcein imaging. Interestingly, RBC Ca^2+^ uptake via piezo1 stimulation led to less RBC death, hemolysis, and cell debris formation than Ca^2+^ ionophore treatment. These results provide evidence that piezo1 activates RBC mechanotransduction pathways distinct from Ca^2+^ ionophore treatment.

EV production methods can profoundly impact EV yield and contents; thus, it is essential to fully characterize how production conditions impact EVs.^29^ RBC-EVs are currently generated *in vitro* anywhere from 2 to 48 hours in the literature,^5,10,30^ or multiple weeks for RBC-EVs generated during storage.^10,30^ RBC stimulation and treatment methods are important considering that the RBCs will eventually consume available glucose, leading to adenosine triphosphate (ATP) depletion that disrupts membrane integrity leading to RBC-EV release.^10,31^ In this study, we found that 6% hematocrit treated with 10 *μ*M yoda1 for as low as 30 minutes minimized hemolysis and maximized EV concentration and purity. We isolated ~10^11^ RBC-EVs/mL from as little as 1mL RBCs, which is sufficient for biological studies given that the total circulating EV concentration is ~10^10^ particles/mL.^21^ Interestingly, we found that 24-hour yoda1 treatment generated RBC-EVs with three distinct size peaks compared to only one peak for RBC-EVs generated under 2 hours. Piezo1 stimulation increases ATP release from human RBCs.^32^ Therefore, piezo1 stimulation may accelerate ATP depletion, causing contamination by EV-subpopulations that mimic storage lesions at longer yoda1 treatment times.

EV isolation methods can similarly impact EV yield and purity. Therefore, we examined how EV isolation via membrane-based affinity, ultrafiltration, and ultracentrifugation affected RBC-EV yield and purity. We found that ultracentrifugation provided the greatest and most consistent piezo1 RBC-EV yield but was the most time-consuming (~3 hours) method and unable to pellet low RBC-EV concentrations generated during needle shearing. Ultrafiltration using Amicon Ultra-15, however, was the more versatile RBC-EV isolation method, requiring a readily available benchtop centrifuge that was able to concentrate shear stress and piezo1 RBC-EVs within 1 hour. However, we found that Amicon Ultra-15 led to less consistent RBC-EV preparations, as there was greater variability in RBC-EV yield than ultracentrifugation. There were no significant differences in protein, protein aggregate, and platelet-EV contamination in RBC-EVs isolated using ultracentrifugation and Amicon Ultra-15. ExoEasy yielded lower RBC-EVs concentrations than Amicon Ultra-15 (*p*>0.05) and ultracentrifugation (*p*<0.05) isolation methods. This low RBC-EV yield caused difficulty when characterizing RBC-EV using Western blot and visualizing EVs using TEM. We also found that piezo1 RBC-EVs isolated using exoEasy contained more protein, protein aggregate, and platelet EVs contamination than RBC-EVs isolated using ultracentrifugation or Amicon Ultra-15.

Pressure myography was used to highlight the importance of EV purity for biological studies. Interestingly, we also found that RBC-EVs isolated using membrane-based affinity caused rapid mouse carotid artery vasodilation, suggesting that RBC piezo1 stimulation may release atheroprotective biomolecules. However, mouse carotid arteries treated with piezo1 RBC-EVs isolated by ultrafiltration and ultracentrifugation, which were more pure, did not vasodilate. Thus, it is likely that particles isolated using membrane-based affinity contained co-isolated protein contaminants that triggered vasodilation. The importance of EV purity is also highlighted by Forteza-Genestrla *et al.* They found that mesenchymal stromal cell EVs isolated using ultracentrifugation (120,000xg for 18 hours) increased metalloproteinases and decorin and decreased collagen and aggrecan mRNA levels in mouse chondrogenic cells.^33^ These pathological effects, however, were mitigated when chondrogenic cells were treated with EVs isolated with ultracentrifugation and purified using SEC.^33^ These data highlight the need to characterize the impact of both RBC-EVs and non-EV contamination in biological studies.

We used SEC in an attempt to further purify our RBC-EVs. In this study, however, SEC did not significantly diminish protein and platelet EV contamination. We found that SEC may decrease protein aggregates, but these results were variable and not statistically significant for RBC-EVs purified after ultrafiltration and ultracentrifugation. Furthermore, RBCs treated with Ca^2+^ ionophore generated greater hemoglobin contamination via hemolysis than yoda1 treatment. However, there were no significant differences in concentration and protein contamination between piezo1 and Ca^2+^ ionophore RBC-EVs, suggesting that ultracentrifugation is able to sufficiently separate hemoglobin contaminants from RBC-EVs generated *in vitro.* We have previously found SEC as an effective method in separating EVs from non-EV contaminants (**Supplementary Fig. 5**). The discrepancy between current and previous study may be caused by how the RBCs are isolated prior to yoda1 treatment. In our previous studies, RBCs did not thoroughly remove the buffy coat, as our objective was to establish a RBC-EV isolation protocol. In the current study, we thoroughly washed the RBCs three times in PBS to minimize platelet EV contamination and obtain the purest RBC-EV prep possible (**Supplementary Fig. 6**). Therefore, RBC-EV purity may be affected by how well the RBCs are isolated from other circulating cell sub-populations and plasma protein contaminates.

We compared RBC-EV purity measurements obtained in this study to literature purity values. Previous studies generally report particle/total protein ratios of 10^7^ to 10^8^ particles/*μ*g when isolating plasma or serum EVs.^34,35^ In this study, RBC-EVs isolated using ultracentrifugation contained particle/total protein ratios of ~10^9^ particles/*μ*g, 10x higher than EVs isolated from circulation. Webber *et al.* proposed particle/total protein criteria to define EV purity, suggesting that ratios > 3×10^10^ particles/*μ*g equate to high EV purity, ratios from 2×10^9^ to 2×10^10^ particles/*μ*g equate to low purity, and any ratio below 1.5×10^9^ particles/*μ*g to be considered impure. However, as noted by the authors, their particles/*μ*g calculations assumed that all detected particles were vesicles and thus do not account for protein aggregates and other contaminates that may overestimate the true number of vesicles isolated. Additionally, particles/*μ*g purity measurements make the assumption that all EV subpopulations contain similar protein concentrations. In reality, EV protein concentration likely depends on the originating cell type and conditions in which EV shedding is stimulated, highlighting the importance of using multiple purity metrics to gain a more holistic understanding of EV contamination.

## Conclusions

We now provide methods to produce RBC-EV via RBC piezo1 stimulation, as well as isolate, verify, and characterize these RBC-EV. Piezo1 stimulation of 6% hematocrit with 10 *μ*M yoda1 for 30 minutes preserved RBC viability, induced less hemolysis, and produced enough RBC-EVs for mechanistic studies from just 1 mL of blood. Ultracentrifugation resulted in the greatest RBC-EV yield and purity, but ultrafiltration required the least time and still provided sufficient RBC-EV yield and purity. This piezo1 RBC-EV production and isolation protocol will help standardize biological studies to show how RBC-EVs generated via physiological shear stress affect vascular health.

## Methods

### Human Participants

All procedures and documents were approved by the University of Maryland Institutional Review Board (IRB Net ID: 1385293-7) and conformed to the Declaration of Helsinki. Healthy men and women (20 to 31 years old) were recruited for up to eight blood draws. All participants provided written informed consent and completed a health history questionnaire. Exclusion criteria included any of the following: current or prior metabolic or cardiovascular disease, inflammatory conditions, cancer, current or previous smoker, use of potential study-confounding medications, anemia, or body weight <110 pounds.

### Blood Collection

Participants reported to the laboratory in the morning following an overnight fast of ≥10 hours. Prior to each blood draw, participants also refrained from using alcohol, non-steroidal anti-inflammatory drugs (NSAIDS), and medications for 24 hours, as well as participating in physical exercise for at least 16 hours. Participants were allowed to continue taking regular oral contraceptives. Additionally, all participants reported not being sick for at least one week prior to blood draws. After a seated rest of ~5 minutes, 20-30 mL of blood was drawn from an antecubital vein into tubes containing acid citrate dextrose (ACD) as an anticoagulant. Blood samples were immediately placed on ice until RBC isolation (usually <1 hour)

### RBC Isolation

Blood was pooled from two male and two female subjects and centrifuged at 1500xg for 5 minutes. Plasma was then removed, and RBCs were washed four times with phosphate buffered saline (PBS) to remove the buffy coat (white blood cells and platelets; **Supplementary Fig. 6**). RBC were resuspended in PBS supplemented with 99 mg/dL glucose and 1 mM CaCl_2_ at pH 7.4.

### RBC Viability Assay

To ensure that RBC were viable before and after EV generation, RBC (5 million cells/mL) were incubated with 2 *μ*g/mL calcein AM (Thermo Fisher Scientific) for 15 minutes at 37°C. RBC were then streaked onto a glass slide, sandwiched with a glass coverslip, and imaged at 20x on a Revolve fluorescence microscope (Echo). RBC viability was determined based on co-localization of calcein AM and brightfield cells in ImageJ. Percent viability was calculated as the sum of all viable RBCs divided by the sum of all particles.

### Hemolysis Measurement

Hemolysis was calculated from the hemoglobin absorbance of 200 *μ*L RBC supernatant at 541 nm using a Spark multimode microplate reader (Tecan, Switzerland).^24,36,37^ Absorbance was used to calculate hemolysis as:^24,37^

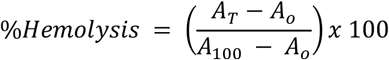

Where A_0_ is the supernatant absorbance from RBCs in PBS, A_100_ is the supernatant absorbance from RBCs in deionized water (100% cell lysis), and A_T_ is the supernatant absorbance from RBCs treated with DMSO, yoda1, or calcium ionophore.

### RBC-EV Production

RBC-EVs were mechanically stimulated by shearing RBCs (6% hematocrit) through a 2 inch, 25G needle 15 times. RBCs not sheared served as controls. We calculated wall shear stress (*τ*) assuming 6% hematocrit viscosity 1.5×10^3^ Pa•s, average volumetric flow rate 8.3m L/min, 25G needle inner diameter 0.280mm.

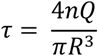

Shear stress-induced RBC-EVs were then concentrated from 50 mL to 400 *μ*L using Amicon Ultra-15 centrifugal filter units (UFC910024, Millipore Sigma; 100 kDa molecular weight cut off). RBC-EVs were also created by treating RBCs with piezo1-agonist yoda1, Ca^2+^ ionophore, or 0.28% DMSO (vehicle control) in a 5% CO_2_, 37°C incubator. The RBC treatment protocol was tuned to improve piezo1 RBC-EV yield and purity using 1, 3, 6, 18% hematocrit, 0.5, 1, 2, and 24 hour yoda1 treatment times, and 1, 10, 100 *μ*M yoda1 concentration. Ca^2+^ ionophore RBC-EVs were used as a gold-standard comparison with 6% hematocrit, 1 or 24 hour treatment times, and 10 or 100 *μ*M Ca^2+^ ionophore concentration. All RBC-EVs were stored at −80°C until needed for experiments.

### RBC-EV Isolation

Plasma EVs were isolated from healthy donors using Total Exosome Isolation Kit (4484450, Invitrogen) according to the manufacturer’s protocol. RBC-EVs generated *in vitro* were isolated using membrane-based affinity binding, ultrafiltration, or ultracentrifugation (**Table 1**). We used the exoEasy Maxi Kit as our representative membrane-based affinity binding method and Amicon Ultra-15 centrifugal units as our representative ultrafiltration method. For all RBC-EV isolation methods, RBC and cell debris were first removed by differential centrifugation at 600xg for 20 minutes, 1600xg for 15 minutes, and 3250xg for 15 minutes at room temperature, followed by 10,000xg for 30 minutes at 4°C. The RBC supernatant was then passed through a 0.22 *μ*M PES syringe filter (371-2115-OEM; Foxx Life Sciences) for ultracentrifugation and Amicon Ultra-15 isolations and 0.8 *μ*M surfactant-free cellulose acetate syringe filter (16592, Sartorius) for exoEasy isolations, as recommended by the vendor. For membrane-based affinity binding, 8 mL RBC supernatant and 8 mL XBP buffer were mixed and passed through the exoEasy affinity spin column to bind RBC-EVs to the membrane. RBC-EVs were then eluted using 500 *μ*L XE buffer, as per manufacturer instructions. For ultrafiltration, 12 mL RBC supernatant was passed through an Amicon Ultra-15 centrifugal filter at 3260xg for 15 minutes. A second Amicon Ultra-15 centrifuge spin was used to wash and concentrate RBC-EVs in 2xPBS. For ultracentrifugation, 35 mL RBC supernatant was centrifuged at 100,000xg for 70 minutes at 4°C. The RBC-EV pellet was washed in PBS, centrifuged again at 100,000xg for 70 minutes, and finally resuspended in 400 *μ*L 2X PBS. RBC-EV aliquots were frozen at −80°C until use; therefore, all RBC-EVs in this study experienced one freeze thaw cycle.

**Table 1:** RBC-EV Isolation Protocol Summary

|  | Ultracentrifugation |  | Amicon Ultra-15 |  | ExoEasy Maxi Kit |  |
| --- | --- | --- | --- | --- | --- | --- |
| Starting Volume | 35 mL |  | 12 mL |  | 8 mL |  |
| Cell Debris Removal | 600xg for 20 minutes |  |  |  |  |  |
|  | 1600xg for 15 minutes |  |  |  |  |  |
|  | 3260xg for 15 minutes |  |  |  |  |  |
|  | 10,000xg for 30 minutes |  |  |  |  |  |
| | Pass supernatant through 0.22 $\mu\text{m}$ filter | | Pass supernatant through 0.22 $\mu\text{m}$ filter | | Pass supernatant through 0.8 $\mu\text{m}$ filter | |
| EV Isolation | 100,000xg for 70 min at 4°C |  | 3,820xg for 15 min at 25°C |  | RBC-EV Sample + Buffer XBP 500xg for 1 min |  |
|  | 100,000xg for 70 min at 4°C |  | 3,820xg for 15 min at 25°C |  | Buffer XWP wash 5000xg for 5 min |  |
|  |  |  |  |  | Buffer XE elution 500xg for 5 min |  |
| Purification | - SEC | + SEC | - SEC | + SEC | - SEC | + SEC |

### Size Exclusion Chromatography (SEC)

A subset of RBC-EVs isolated using membrane-based affinity binding, ultrafiltration, or ultracentrifugation were further purified using SEC. 500 *μ*L RBC-EV sample was added to SEC columns (qEV original, IZON) and eluted with 2X PBS. Six 500 *μ*L eluate fractions were collected. Protein concentration was measured to identify fractions containing RBC-EVs by spectrophotometrically reading absorbance at 280 nm (NanoDrop 2000c, Thermo Fisher Scientific).

### EV Size and Concentration

Tunable resistive pulse sensing (TRPS) was used to quantify RBC-EV concentration and size (50-330nm) using a qNano Gold (IZON Science) with NP100 nanopore membranes. RBC-EV samples were first filtered through 0.22 *μ*m ultrafree-MC tubes (UFC30GV, Millipore Sigma) to remove large particles that could clog the nanopore. TRPS voltage was set between 0.5-0.7 V to achieve a stable 130 nA current. Particles were counted when root mean square (RMS) noise was below 10 pA and particle rate was linear. At least 500 particles were counted at particle rates between 200-1500 particles/minute and 5 or 10mBar pressure. TRPS was calibrated using CPC100 beads (mean diameter: 100nm).

### Transmission Electron Microscopy (TEM)

TEM was used to confirm RBC-EV size and visualize RBC-EV shape. We applied 5 *μ*L RBC-EV sample to a 200 mesh copper grid coated with formvar carbon film (FCF200-Cu, Electron Microscopy Sciences) for 30 seconds. The copper grid was washed with PBS and stained using 5 *μ*L 2% uranyl acetate for 30 seconds. TEM images were acquired using a Hitachi HT7700 Transmission Electron Microscope.

### Western Blot

Western blot was performed to detect specific RBC and RBC-EV proteins. RBCs were washed with ice-cold PBS and then lysed with RIPA buffer (Thermo Fisher Scientific) that contained Halt Protease Inhibitor Cocktail (Thermo Fisher Scientific) and EDTA (Thermo Fisher Scientific). Lysates were centrifuged for 15 min at 4°C to remove cellular debris. Lysate protein was quantified and normalized using a BCA assay (Thermo Fisher Scientific). We loaded 200 *μ*g RBC protein lysate into each well of a NuPAGE 4-12% Bis-Tris gel (Thermo Fisher Scientific) for protein separation by SDS-PAGE. RBC-EVs were lysed by sonicating 3 times for 15 seconds each. We then loaded 29 *μ*L of lysed RBC-EVs in NuPAGE 4-12% Bis-Tris protein gels (Thermo Fisher Scientific). Separated proteins were transferred onto polyvinylidene difluoride (PVDF) membranes using an iBlot 2 (Thermo Fisher Scientific) semi-dry transfer system. PVDF membranes were blocked in 5% bovine serum albumin (BSA) in TBS-Tween (TBS-T) for one hour at room temperature. Blots were then incubated with primary antibodies in 1% BSA in TBS-T overnight at 4°C. Primary antibodies included ALIX (1:500; sc-53538), TSG101 (1:500; sc-7964), CD9 (1:500; sc-13118), CD63 (1:500; sc-5275), calnexin (1:500; sc-23954), stomatin (1:500; sc-376869), CD52 (1:500; sc-51560), p-eNOS (1:500; 9571S), eNOS (1:500; 9572S), and GAPDH (1:1000; 2118S; all from Cell Signaling Technology). Blots were then incubated with the appropriate mouse or rabbit horseradish peroxidase conjugated secondary antibody (Thermo Fisher Scientific) for two hours at room temperature. Protein bands were detected using a chemiluminescence kit (SuperSignal West Pico PLUS, Thermo Fisher Scientific) and imaged immediately using an Alpha Innotech Fluorchem Imager. ImageJ (NIH) was used to quantify band intensity, which was normalized to GAPDH or β-actin.

### Flow Cytometry

Flow cytometry was used to classify the cellular origin of isolated EVs based on surface markers. EVs were labeled for 45 minutes at room temperature in the dark with Alexa Fluor 647 conjugated anti-CD63 (Clone: H5C6; 353015), BV421 conjugated anti-CD41 (Clone: HIP8; 303729), PE dazzle 594 conjugated anti-CD235a (Clone: HI264; 349119) and BV605 conjugated anti-CD31 (Clone: WM59; 303121; all from Biolegend). EVs were then resuspended in 500 *μ*L particle free PBS and analyzed using a BD FACSAria Flow cytometer (Becton Dickenson, Vancouver BC, Canada). Data were analyzed using FlowJo 10.8.0 software (Becton Dickenson, Vancouver BC, Canada). Buffer and unlabeled EVs were the negative controls. Forward scatter (FSC) and side scatter (SSC) were set to gate the EV population. Each marker expression was represented by histogram and normalized to mode. 200 nm calibration beads were used for FSC and SSC parameters.

### Protein Contamination

RBC-EV protein contamination was measured using the particle/total protein ratio (particles/*μ*g protein). RBC-EV concentration was calculating using TRPS. Sample protein concentration was measuring using a Bicinchoninic acid (BCA) protein assay kit (Thermo Fisher Scientific, MA, USA), with bovine serum albumin standards ranging from 20-2000 *μ*g/mL.

### Protein Aggregates

Protein aggregates (non-EV contaminants) were quantified using the EV/Non-EV particle rate ratio. Non-EV samples were created by vortexing samples in 1% Triton-X 100 surfactant every 15 minutes for 1 hour to lyse EVs.^38^ Particle rate was then measured in lysed and intact RBC-EV samples by counting particles via TRPS over 1 minute. Particle rate was used as a surrogate for concentration because a particle rate of 100 particles/min is needed to accurately quantify size and concentration, and lysed samples never reached this particle rate.

### Animals

The animal protocol was approved by the University of Maryland Institutional Animal Care and Use Committee (IRBNet ID: 1664120-1). 12-week old male C57Bl/6 mice were acquired from Jackson Laboratory (Bar Harbor, ME) and maintained on a normal chow diet *ad libitum*. Animals were acclimated for at least three days in the University of Maryland vivarium prior to euthanasia by exsanguination. The carotid arteries were quickly excised and placed in cold HEPES physiological saline solution (HEPES-PSS) and used within 24 hours. HEPES-PSS (6.96g NaCl, 0.35g KCl, 0.29g MgSO_4_*7H_2_O, 0.16g KH_2_PO_4_, 0.31g CaCl_2_*2H_2_O, 1.25g NaHCO_3_, 2.38g HEPES, 0.99g glucose, 2mL 500x EDTA and pH 7.4) was made fresh on the experiment day.

### Pressure Myography

Carotid arteries were cannulated with stainless steel micropipettes in a pressure myograph chamber (114P; DMT) containing cold HEPES-PSS. The myograph chamber was then slowly warmed to 37°C and aerated with carbogen (5% CO_2_, 95% O_2_). Arterial pressure was increased in 10 mmHg increments, every 5 minutes, up to 40 mmHg. Carotid arteries were then allowed to equilibrate at 50 mmHg for an additional 15 minutes. Vessel viability was confirmed by superfusing arteries with high potassium PSS (KPSS; 3.69g NaCl, 18.6g KCl, 0.36g MgSO_4_*7H_2_O, 0.41g KH_2_PO_4_, 0.46g CaCl_2_*2H_2_O, 0.21g NaHCO_3_, 0.18g glucose, 95 *μ*L 500x EDTA) for 5 minutes. Carotid arteries were considered viable if the vessel constricted at least 10%. KPSS tests were repeated two times. The carotid arteries were then washed with HEPES-PSS and submaximally constricted 10-15% using 10^−6^ M phenylephrine (207240100; Acros Organics). Vasodilation was measured by treating carotid arteries for five minutes with RBC-EVs isolated using exoEasy, Amicon Ultra-15, and ultracentrifugation. Finally, endothelial dependent vasodilation was measured in the same carotid arteries using an acetylcholine dose response ranging from 10^−9^ to 10^−5^ M acetylcholine. Vasodilation was measured using the maximum steady-state diameter (D_a_), steady-state diameter after phenylephrine preconstruction (D_p_), and maximal steady-state baseline vessel diameter (D_m_).

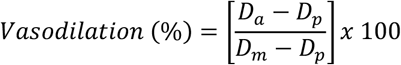

### Hemoglobin ELISA

Hemoglobin concentration in RBC-EVs was measured using a hemoglobin ELISA (Thermo Fisher Scientific) as per manufacturer protocol. RBC-EV were diluted 250,000x with the appropriate assay diluents. Samples were assayed in duplicate. Immediately after the addition of stop solution, absorbance was measured at 450nm via spectrophotometer (Tekan Spark Multimode Microplate Reader).

## Acknowledgments

Funding support was provided to G.S.S through the University of Maryland Presidential Postdoctoral Fellowship and the American Heart Association Postdoctoral Fellowship (916512), C.M.W. through the National Science Foundation Graduate Research Fellowship Program (DGE 1840340), R.M.S by a National Institutes of Health Postdoctoral Institutional Training Grant (5T32HL007698-26) and an American Heart Association Postdoctoral Fellowship (915916), and P.B through National Institute of Health grants (R01HL156526, R01HL159862, R01HL158076, R01HL161004, R01HL162120), and A.M.C through a National Institute of Health grant (R01HL140239–0).

## Contributions

G.S.S. planned, performed, and coordinated all experiments, wrote the manuscript, and created the figures. C.M.W. assisted RBC-EV isolation using ultracentrifugation, performed all RBC-EV isolations and SEC purifications using Amicon Ultra-15 centrifuge units, and quantified RBC viability using Calcein imaging. R.M.S. conducted all blood draws, performed all RBC-EV isolation and SEC purifications using exoEasy, and completed hemoglobin ELISA. S.S. and K.T. and performed flow cytometry experiments and helped troubleshoot RBC-EV verification. M.P. helped establish RBC-EV isolation protocols by verifying EV markers using Western blot. P.B. and A.D. provided feedback on experiments, manuscript, and figures. A.M.C. oversaw all experiments, as well as the preparation of manuscript and figures.

