## Supplementary Figures for "Mechanical Stimuli such as Shear Stress and Piezo1 Stimulation Generate Red Blood Cell Extracellular Vesicles"

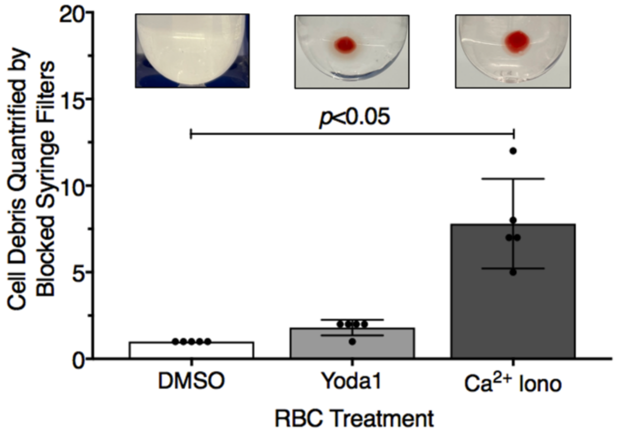


**Supplementary Fig. 1 | $Ca^{2+}$ ionophore treatment generates more cell debris than untreated and yoda1 treated RBCs.** RBC debris after DMSO, yoda1, and $Ca^{2+}$ ionophore treatment, as quantified by the number of 100% blocked syringe filters after 10,000xg centrifugation. Inset show representative image of RBC debris pellet after 10,000xg centrifugation.

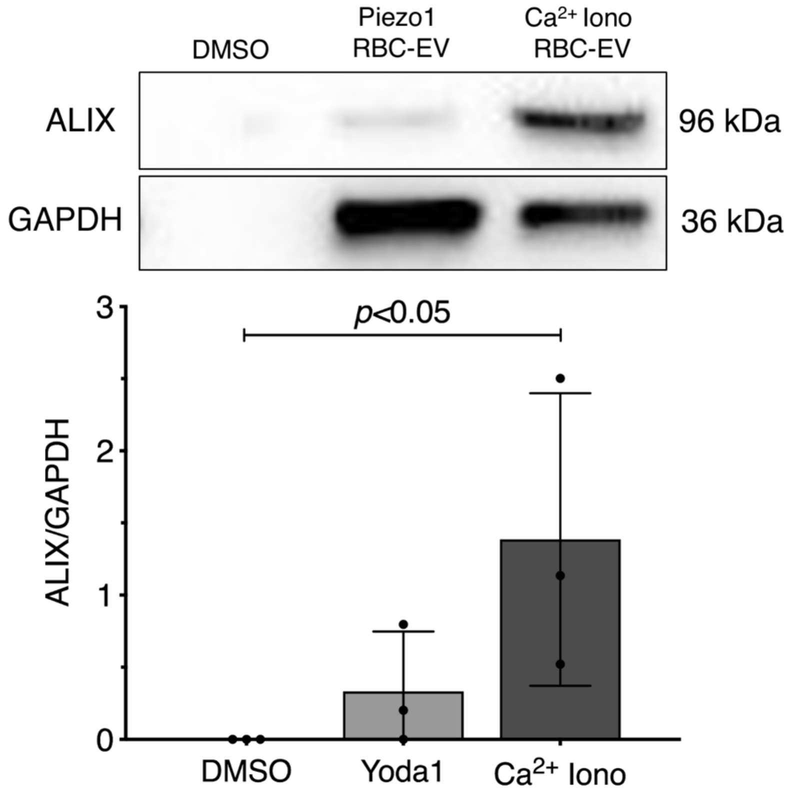


**Supplementary Fig 2. | $Ca^{2+}$ ionophore RBC-EVs contain greater ALIX than piezo1 RBC-EVs.** Western blots (9.75 µL sample/well) of cytoplasmic EV markers ALIX and housekeeper GAPDH in particles released after DMSO treatment and in Piezo1 and $Ca^{2+}$ ionophore RBC-EVs. Western blot shows one representative comparison.

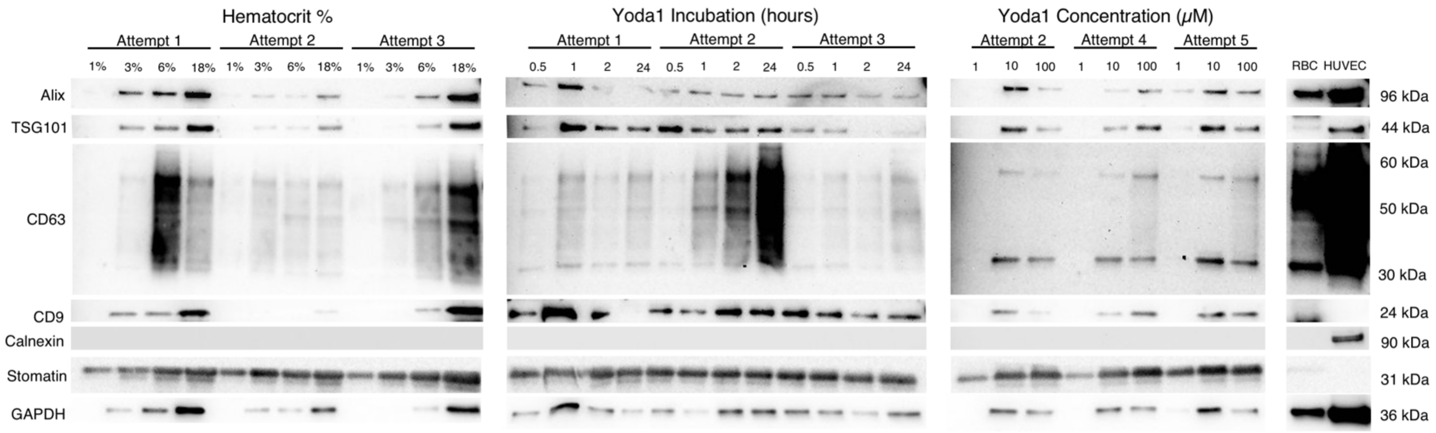


**Supplementary Fig. 3 | RBC-EV markers vary depending on treatment conditions and biological replicate.** Western blots (9.75 µL sample/well) for RBC-EV, RBC-specific, and apoptotic body markers in **a.** hematocrit dose response**, b.** yoda1 treatment time, **c.** yoda1 dose response experiments**,** and **d.** RBCs and HUVEC lysates.

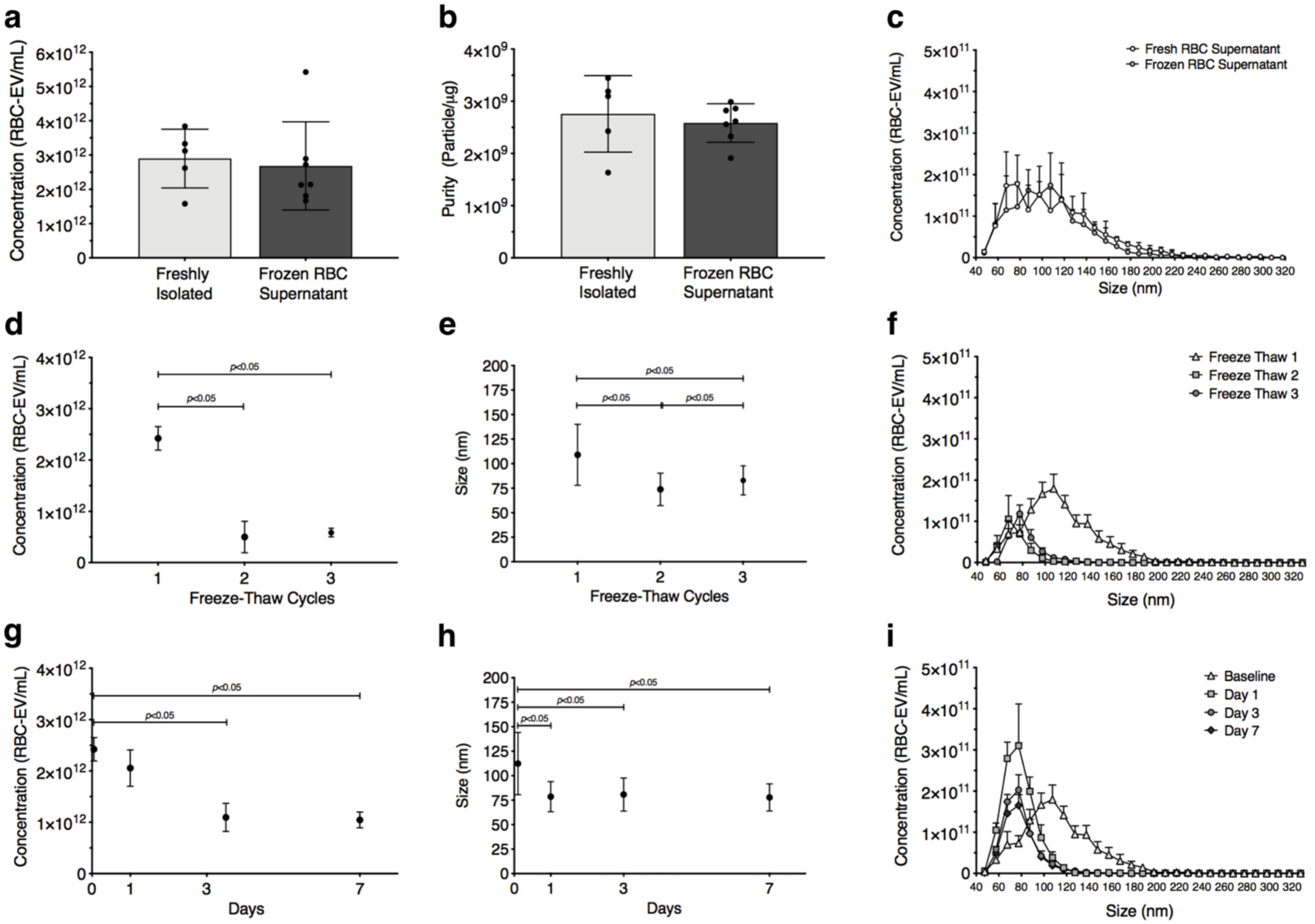


**Supplementary Fig. 4 | RBC-EV concentration and size changes with storage conditions. a.** Concentration, **b.** purity, and **c.** size histogram of RBC-EVs isolated using fresh RBC supernatant and frozen RBC supernatant. **d.** Concentration, **e.** size, and **f.** size histogram of RBC-EVs undergone up to 3 freeze thaw cycles. **g.** Concentration, **h.** size, and **i.** size histogram of RBC-EVs stored in 4°C for up to 7 days. Size and concentration were measured using TRPS and purity was measured by calculating particle/total protein ratio.

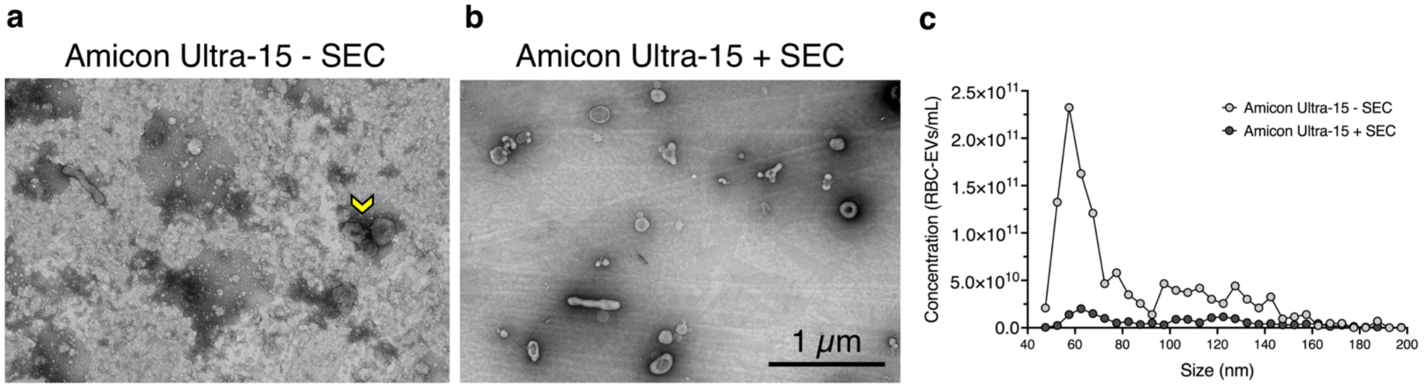


**Supplementary Fig. 5 | SEC improves RBC-EV purity in highly contaminated samples.** TEM of RBC-EVs isolated using Amicon Ultra-15 centrifuge units **a.** with and **b**. without SEC purification. Yellow arrow highlights RBC-EVs in contaminated samples. **c.** TRPS size histogram of RBC-EVs isolated using Amicon Ultra-15 and then purified with or without SEC.

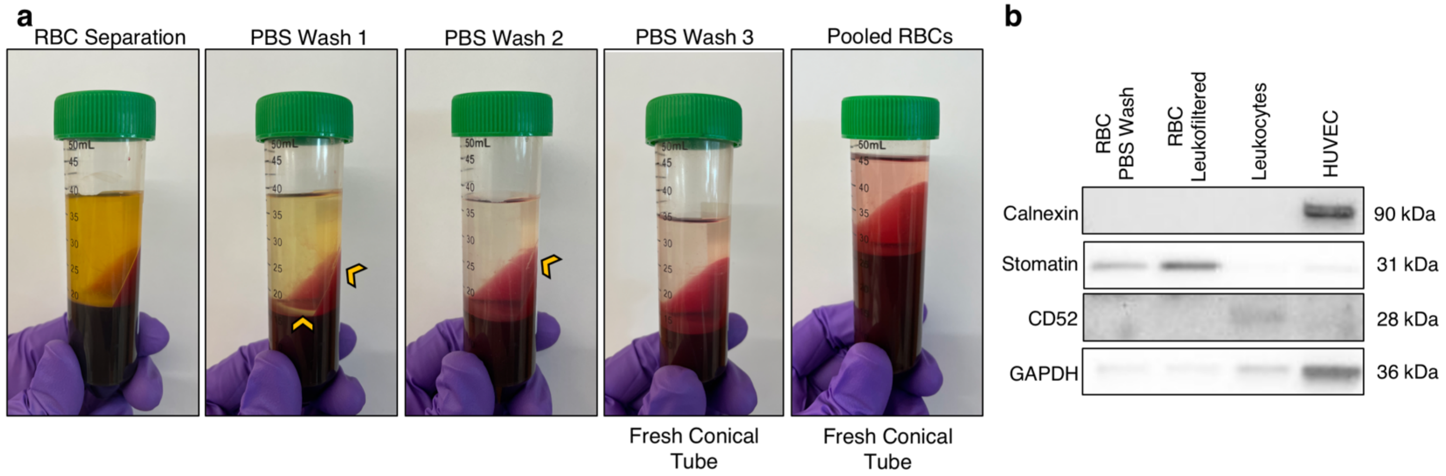


**Supplementary Fig. 6 | Triple washing RBCs in PBS sufficiently removed leukocytes and other large cells. a.** Images illustrating how RBCs were washed and isolated from plasma and buffy coat. RBCs were washed until contamination (yellow arrow) was removed. **b.** Western blot showing that RBCs washed in PBS was able to remove leukocytes (CD52) and other cell types (calnexin) as well as leukofiltered RBCs.
